# A direct comparison of genome alignment and transcriptome pseudoalignment

**DOI:** 10.1101/444620

**Authors:** Lynn Yi, Lauren Liu, Páll Melsted, Lior Pachter

## Abstract

**Motivation:** Genome alignment of reads is the first step of most genome analysis workflows. In the case of RNA-Seq, transcriptome pseudoalignment of reads is a fast alternative to genome alignment, but the different “coordinate systems” of the genome and transcriptome have made it difficult to perform direct comparisons between the approaches.

**Results:** We have developed tools for converting genome alignments to transcriptome pseudoalignments, and conversely, for projecting transcriptome pseudoalignments to genome alignments. Using these tools, we performed a direct comparison of genome alignment with transcriptome pseudoalignment. We find that both approaches produce similar quantifications. This means that for many applications genome alignment and transcriptome pseudoalignment are interchangeable.

**Availability and Implementation:** bam2tcc is a C++14 software for converting alignments in SAM/BAM format to transcript compatibility counts (TCCs) and is available at https://github.com/pachterlab/bam2tcc. kallisto genomebam is a user option of kallisto that outputs a sorted BAM file in genome coordinates as part of transcriptome pseudoalignment. The feature has been released with kallisto v0.44.0, and is available at https://pachterlab.github.io/kallisto/.

**Supplementary Material:** N/A

**Contact:** Lior Pachter

## Introduction

Read alignment programs are used to locate the genome coordinates from which a sequenced read could originate (e.g.,Langmead and Salzberg, 2012). In the case of RNA-Seq, the sequenced reads correspond to cDNA that have been reverse transcribed from mRNA. Because splicing occurs as part of post-transcriptional processing, the alignments to the genome may span multiple exons and skip the introns between the exon-exon junctions. The task of aligning RNA-Seq reads to the genome in a way that is robust to splicing is known as “genome spliced alignment.” There are several programs that perform this task, such as TopHat/TopHat2 (Trapnell *et al.*, 2009; Kim *et al.*, 2013), STAR (Dobin *et al.*, 2013), and HISAT/HISAT2 (Kim *et al.*, 2015). When spliced alignment is used for RNA-Seq, subsequent analysis is required to assign reads to genes (e.g.,Liao *et al.*, 2014) or to transcripts (e.g.,Trapnell *et al.*, 2010) as part of quantification.

An alternative to align reads to the genome is to align reads directly to the transcriptome. The transcriptome is defined as the set of sequences corresponding to mature mRNA after post-transcriptional processing. Methods such as eXpress (Roberts and Pachter, 2013) and RSEM (Li and Dewey, 2011) use transcriptome alignments for read assignment under a quantification model. In previous benchmarks of RNA-Seq quantification methods, it has been unclear whether performance improvements from transcriptome alignment are due to the mode of alignment or to a different quantification model (Teng *et al.*, 2016).

In 2016, Bray *et al*. introduced the concept of pseudoalignment to the transcriptome, which, rather than a performing a full alignment, records information about the set of transcripts a read is compatible with. Specifically, in pseudoalignment, reads are assigned to these sets of transcripts, i.e. “equivalence classes” of transcripts (Nicolae *et al.*, 2011). Transcript compatibility counts (TCCs) constitute the number of reads within the equivalence classes and serve as sufficient statistics for transcript quantification. (See Figure 1a for workflow of pseudoalignment and quantification.) Transcriptome pseudoalignment is orders-of-magnitude faster than traditional genome alignment (Bray *et al.*, 2016; Patro *et al.*, 2017), however while pseudoalignment has increased in popularity since its introduction two years ago (Vivian *et al.*, 2017; Lachmann *et al.*, 2018), genome alignment programs are still widely used.

**Figure 1.**
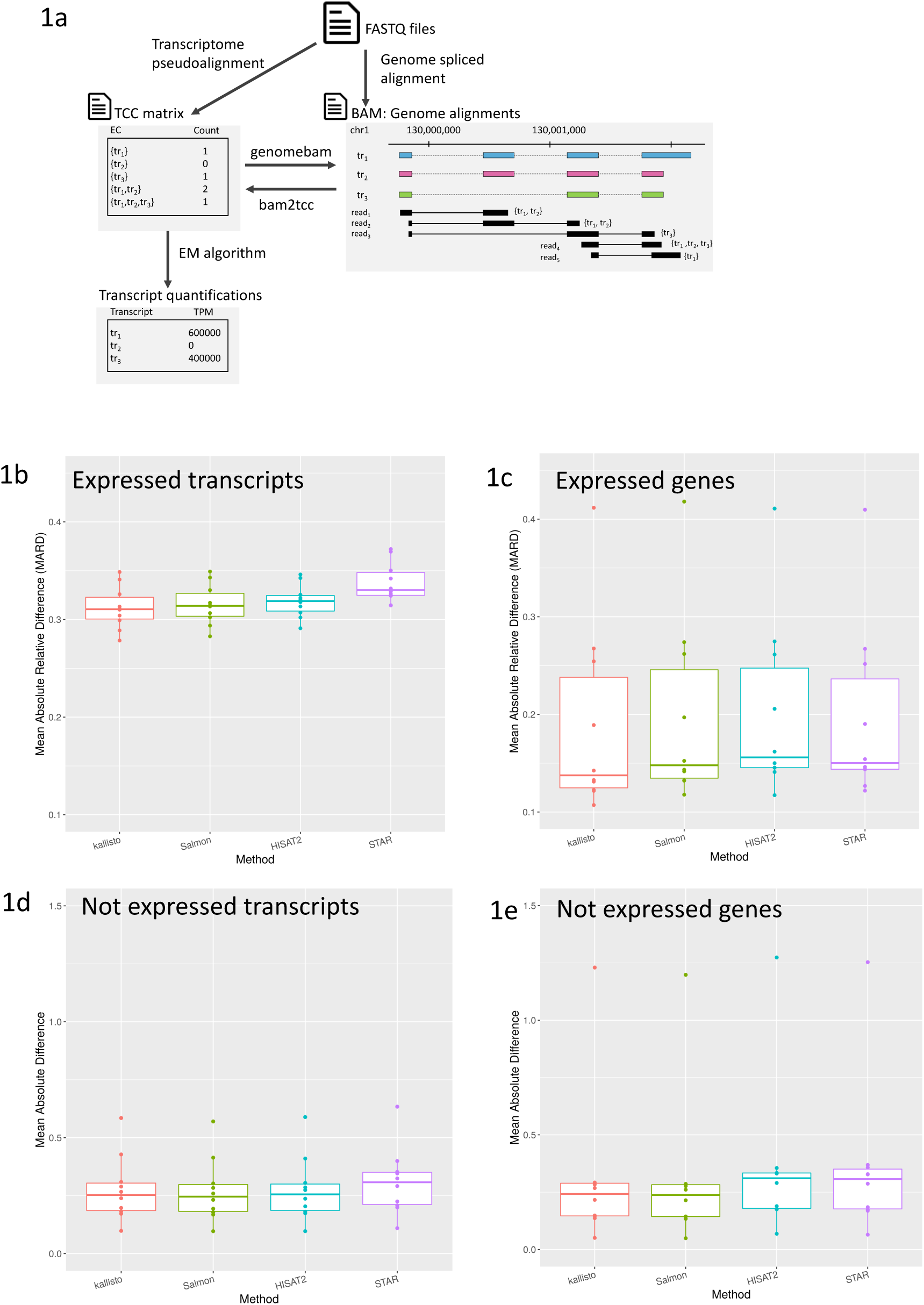
**(a)** We compared genome aligners and pseudoaligners by obtaining transcript compatibility counts from all methods and using kallisto to perform the same EM quantification. bam2tcc was used to convert genome alignments from HISAT and STAR to transcript compatibility counts prior to quantification. We then plotted the mean absolute relative distances (MARDs) across ten simulations for transcripts and genes where the true expression is nonzero **(b-c)** and the mean absolute distance for transcripts and genes where the true expression is zero **(d-e)**.

While there have been comparisons of genome alignment and pseudoalignment methods (Teng *et al.*, 2016; Bray *et al.*, 2016; Patro *et al.*, 2017), the benchmarks have examined only the final quantifications and have not teased apart the algorithmic components. The output of peudoalignment, in the form of TCCs, is conceptually different from genome alignments. To compare genome alignments directly to pseudoalignments, one must perform a conversion of the underlying data models, a task that is considerably more complicated than converting file formats. Furthermore, procedures for quantification after genome alignments are fundamentally different than those that are used with pseudoalignments, making a direct comparison challenging.

## Results

To compare alignment to pseudoalignment methods, we created a tool, bam2tcc, that converts genome alignments in the format of a BAM or SAM file to transcript compatibility counts, the primary output of transcriptome pseudoalignment. We then quantified genome alignments and transcriptome pseudoalignments using the exact same model and method (Figure 1b), thus separating the effects of alignment from those of quantification. We used bam2tcc to convert HISAT2 and STAR genome alignments into transcript compatibility counts, which were then quantified using the expectation maximization (EM) algorithm for a uniform coverage model (Bray *et al.*, 2016). We chose HISAT2 and STAR because of their popularity as well as their accuracy in previous benchmarks (Baruzzo *et al.*, 2017).

We compared the accuracy of the genome spliced alignment programs HISAT2 and STAR and transcriptome pseudoalignment programs kallisto and Salmon on simulations where the true abundances are known. We ran the methods with default parameters and with minimal parameterization. Performance of aligners and pseudoaligners on the simulations were comparable, as demonstrated by their mean absolute relative differences (MARDs) on expressed transcripts. We separately benchmarked accuracy on transcripts that were not expressed in the simulation by examining their distances from zero and taking the mean of this distance across all transcripts. On both measures, across 10 simulated samples, the results of methods were highly concordant with each other (Figure 1b-e), with the exception of STAR. kallisto, Salmon and HISAT2 were more accurate than STAR (Tables 1-4), demonstrating that transcriptome pseudoalignment methods can outperform genome alignment methods.

**Table 1:** Pearson Correlations on Transcript Counts (log(counts+1)), Sim

|  | kallisto | Salmon | HISAT2 | STAR | ground truth |
| --- | --- | --- | --- | --- | --- |
| kallisto | 1 | 0.998 | 0.986 | 0.977 | 0.951 |
| Salmon | 0.998 | 1 | 0.987 | 0.977 | 0.951 |
| HISAT2 | 0.986 | 0.987 | 1 | 0.977 | 0.949 |
| STAR | 0.977 | 0.977 | 0.977 | 1 | 0.941 |
| ground truth | 0.951 | 0.951 | 0.949 | 0.941 | 1 |

**Table 2:** Spearman Correlations on Transcript Counts (log(counts+1)), Sim

|  | kallisto | Salmon | HISAT2 | STAR | ground truth |
| --- | --- | --- | --- | --- | --- |
| kallisto | 1 | 0.991 | 0.950 | 0.920 | 0.840 |
| Salmon | 0.991 | 1 | 0.950 | 0.919 | 0.840 |
| HISAT2 | 0.950 | 0.950 | 1 | 0.922 | 0.842 |
| STAR | 0.920 | 0.919 | 0.922 | 1 | 0.819 |
| ground truth | 0.840 | 0.840 | 0.842 | 0.819 | 1 |

**Table 3:** Pearson Correlations on Gene Counts (log(counts+1)), Sim

|  | kallisto | Salmon | HISAT2 | STAR | ground truth |
| --- | --- | --- | --- | --- | --- |
| kallisto | 1 | 1.000 | 0.997 | 0.996 | 0.992 |
| Salmon | 1.000 | 1 | 0.997 | 0.996 | 0.992 |
| HISAT2 | 0.997 | 0.997 | 1 | 0.997 | 0.989 |
| STAR | 0.996 | 0.996 | 0.997 | 1 | 0.989 |
| ground truth | 0.992 | 0.992 | 0.989 | 0.989 | 1 |

**Table 4:** Spearman Correlations on Gene Counts (log(counts+1)), Sim

|  | kallisto | Salmon | HISAT2 | STAR | ground truth |
| --- | --- | --- | --- | --- | --- |
| kallisto | 1 | 0.998 | 0.987 | 0.985 | 0.982 |
| Salmon | 0.998 | 1 | 0.987 | 0.985 | 0.982 |
| HISAT2 | 0.987 | 0.987 | 1 | 0.986 | 0.975 |
| STAR | 0.985 | 0.985 | 0.986 | 1 | 0.974 |
| ground truth | 0.982 | 0.982 | 0.975 | 0.974 | 1 |

We also compared the results of the four methods on experimental RNA-Seq data from Zika-infected human neuroprogenitor cells (Tang *et al.*, 2016; Yi *et al.*, 2017). Since a simulation cannot capture all sources of variance in an experiment, the inter-method correlations on experimental data were lower than on simulated data. Nonetheless, the cross-method correlations show that the methods still produced concordant quantifications (Tables 5-6).

**Table 5:** Pearson correlation on Transcript Counts (log(counts+1)), Zika

|  | kallisto | Salmon_0.8.2 | Salmon_0.11.2 | HISAT2 | STAR |
| --- | --- | --- | --- | --- | --- |
| kallisto | 1 | 0.998 | 0.966 | 0.936 | 0.934 |
| Salmon_0.8.2 | 0.998 | 1 | 0.966 | 0.936 | 0.934 |
| Salmon_0.11.2 | 0.966 | 0.966 | 1 | 0.970 | 0.966 |
| HISAT2 | 0.936 | 0.936 | 0.970 | 1 | 0.976 |
| STAR | 0.934 | 0.934 | 0.966 | 0.976 | 1 |

**Table 6:** Spearman correlation on Transcript Counts (log(counts)+1), Zika

|  | kallisto | Salmon_0.8.2 | Salmon_0.11.2 | HISAT2 | STAR |
| --- | --- | --- | --- | --- | --- |
| kallisto | 1 | 0.995 | 0.916 | 0.861 | 0.857 |
| Salmon_0.8.2 | 0.995 | 1 | 0.917 | 0.861 | 0.858 |
| Salmon_0.11.2 | 0.916 | 0.917 | 1 | 0.934 | 0.926 |
| HISAT2 | 0.861 | 0.861 | 0.934 | 1 | 0.953 |
| STAR | 0.857 | 0.858 | 0.926 | 0.953 | 1 |

Examining the TCCs derived by each method on this experimental dataset also shows concordance on the level of the TCCs, even prior to quantification. We examined the number of transcripts in each equivalence classes as proxy for the uncertainty in read assignment. The distribution of equivalence class size, defined as the number of transcripts per equivalence class, weighted by the counts for the equivalence class, was similar for all methods (Figure 2a). The weighted mean equivalence class size was similar across all methods and showed that each read on average is compatible with 3.8 ambiguous transcripts (Figure 2b). An examination of the intersections of identical equivalence classes across methods showed that the majority of equivalence class were shared amongst all methods, and that almost all of the reads were in equivalence classes that were common to all methods (Figure 2c).

**Figure 2.**
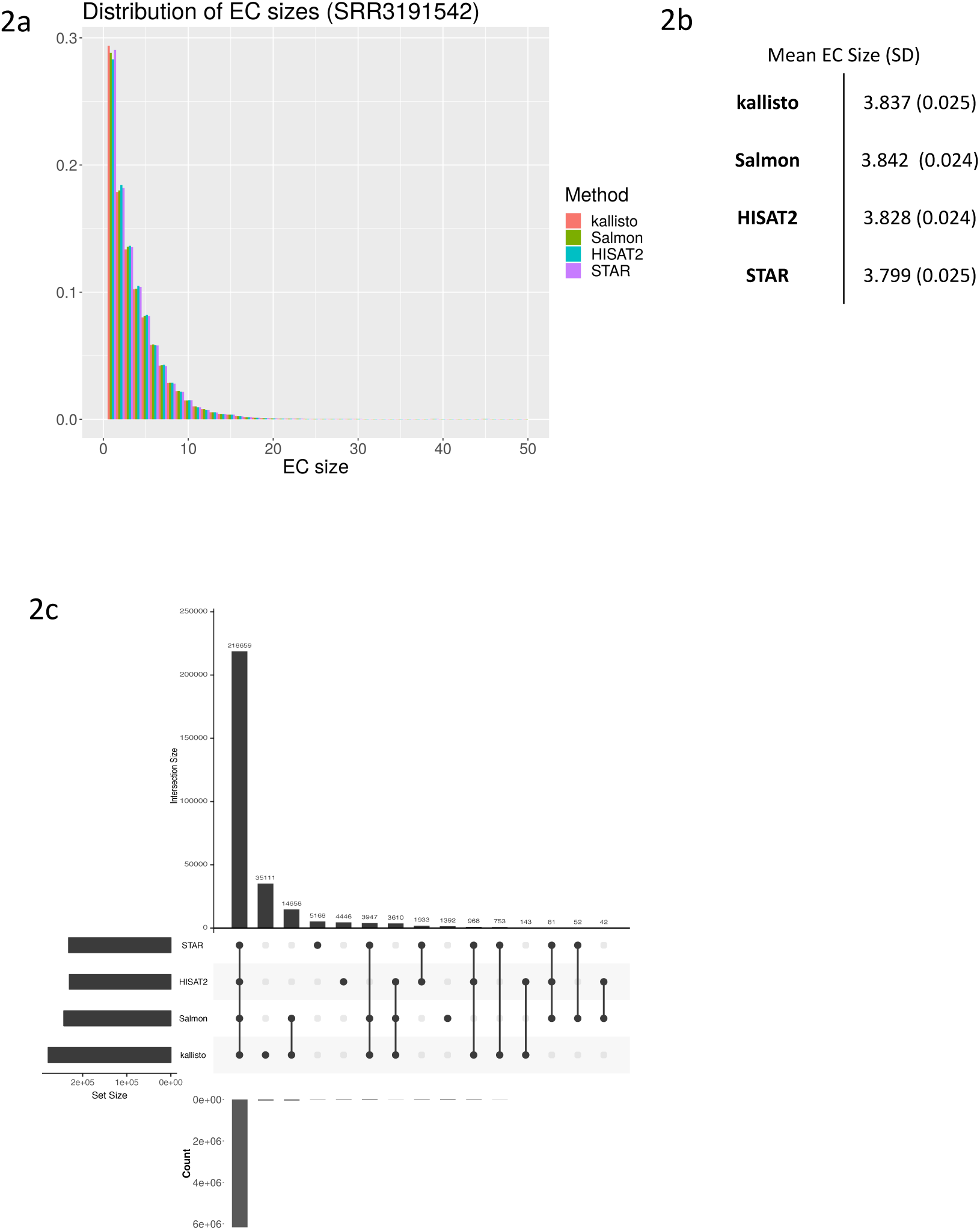
Distribution of equivalence class sizes in a dataset of paired-end RNA-Seq of human neuroprogenitor cells (SRR3191542). The size of an equivalence class is measured as the number of transcripts, weighted by the number of counts in the equivalence class. All methods have similar distributions of equivalence class sizes **(a)**, and furthermore, the methods have comparable mean equivalence class size across the four samples in this dataset **(b)**. The other three samples in the dataset also had similar distributions of equivalence class sizes (data not shown). **(c)** shows the equivalence classes that are shared across methods using an upset plot. The number of shared equivalence classes across the methods are plotted in the top bar graph. The read density in these equivalence classes are plotted in the bottom bar graph, which was calculated as the sum of the counts of the ECs within that intersection.

One feature of genome alignment methods is that the output can be used to produce a visualization of the reads along the genome. Such visualizations are important for quality control and interpretation. To enable the feature with transcriptome pseudoalignment, we developed a tool, *kallisto quant--genomebam*, that generates a BAM file that can be used for visualization, alongside kallisto’s usual quantification. This will allow users to benefit from the speed of pseudoalignment while still being able to visualize the pseudoalignments.

## Discussion

Our analysis is the first direct comparison that specifically examines the differences in alignment compared to pseudoalignment. Whereas previous comparisons confounded alignment/pseudoalignment with quantification, we have controlled for quantification by developing a new tool to convert genome alignments to TCCs. One application of our tool is to convert preexisting alignments into pseudoalignments. Previous work has shown that using TCCs directly for clustering and differential expression is as good as or better than using transcript quantifications (Ntranos *et al.*, 2016; Yi *et al.*, 2018; Ntranos *et al.*, 2018). Direct analysis of TCCs can be advantageous since it does not introduce inferential ambiguity through transcript assignment and is feasible even in the absence of full length sequencing (Ntranos *et al.*, 2016; Yi *et al.*, 2018; Ntranos *et al.*, 2018).

In choosing the benchmarking metrics for our analyses, we separated analysis of expressed transcripts from non-expressed transcripts. This is not typically done but we found that such a separation is important as metrics such as mean absolute difference (MARD) can be biased by zeroes. Because relative differences are more meaningful on expressed transcripts and absolute differences are more meaningful on non-expressed transcripts, we propose that subsequent benchmarks should always separately evaluate the two.

One advantage of performing genome spliced alignment with RNA-Seq reads is that alignments can be readily visualized on browsers (e.g.,Robinson *et al.*, 2011). We provide, for the first time, a tool for visualizing pseudoalignments as projections to the genome. Previously, the pseudoalignment programs RapMap (Srivastava *et al.*, 2016) and kallisto could output SAM formatted alignments, but only with respect to the transcriptome, and were therefore not directly useful for visualization.

Finally, our results demonstrate a practical point for bioinformaticians: for the purpose of transcript quantification, transcriptomic pseudoaligners perform as accurately as aligners. One key advantage of pseudoaligners is speed, and with our new feature, we can support visualization of the pseudoalignments in genomic coordinates. Aside from cases where alignment to noncoding regions is valued (e.g. when transcriptome annotations are incomplete) or where alignments are important for the biology of interest (e.g. for the discovery of novel splice junctions), pseudoalignment should suffice.

## Conclusion

In a first direct comparison between aligners and pseudoaligners, we showed that pseudoaligners are as accurate as genome aligners. We created tool that converts genome alignment in the form of a SAM/BAM into TCCs that can be quantified with kallisto. Furthermore, we implemented a new feature in kallisto for projecting pseudoalignments to the genome, which is output as a BAM file and can be visualized like genome alignments. Our tools place genome alignment and transcriptome pseudoalignment on an equal footing.

## Methods

### bam2tcc

bam2tcc is written in C++14 and uses the SeqAn software library (Reinert *et al.*, 2017) for efficient parsing of BAM and GTF files. bam2tcc requires as inputs a GTF/GFF file for the annotation and a sorted BAM or SAM file of alignments. The output of bam2tcc is a vector of TCCs and a map of ECs to transcripts.

Briefly, bam2tcc first combines the transcript coordinates and the sorted read alignments. For each alignment, and every transcript, it considers whether the alignment is compatible with the transcript based on the exon coordinates of the transcript. An alignment is compatible with a transcript if it starts within an exon of the transcript, ends within an exon of the transcript, and its gaps coincide within the start and end coordinates of all the exons between the start and end exon. For each alignment, the set of transcripts that are compatible with the alignment is its equivalence class. In the case of reads with multiple genome alignments, bam2tcc computes the union of the alignments’ equivalence classes to obtain the equivalence class of the read. For paired-end sequencing, bam2tcc takes the intersection of the equivalence classes corresponding to the two reads to obtain the equivalence class of the pair.

### GenomeBam

kallisto v0.44.0 adds a new option of projecting pseudoalignments of reads to genomic coordinates, where alignments are annotated with the posterior probability of the alignment. To this end, using a user-provided GTF file, kallisto constructs a model of the transcriptome consisting of genes, transcripts and exon coordinates. The reporting of the alignment uses a two-stage process. In the first stage, kallisto performs pseudoalignment and the equivalence class of each read is recorded on disk in a temporary file. Following pseudoalignment, the EM algorithm is run to obtain transcript quantifications. This is the usual workflow of kallisto quantification. In the second stage, with quantification results available, kallisto then loads the temporary file of equivalence classes in conjunction with the reads. For each read, kallisto identifies the first k-mer in the read that maps to the transcripts of the equivalence class with non-zero abundances. Using coordinates of the k-mer within the transcript, kallisto then projects the transcript coordinates to genome coordinates, accounting for exon structure. This subsequent projection is done without additional sequence information beyond the first matching k-mer. The set of genome projections are collapsed, such that a read mapping to multiple transcripts but to a single genomic position has a single alignment record in the BAM file. All multiple genome alignments are reported, but the alignment supported by the highest transcript abundance is reported as the primary alignment. The genome is divided into fixed intervals and each alignment is written to a temporary BAM file on disk corresponding to the interval. After all reads have been processed, each temporary BAM file is sorted and concatenated to a final sorted BAM file. Finally, the sorted BAM file is indexed for fast random access.

### Datasets

We used RSEM v1.3.0 to simulate paired end RNA-Seq samples with uniform coverage. The RSEM model was built using data from single cell RNA-Seq (SMART-Seq) performed on differentiating myoblasts (Trapnell *et al.*, 2014). With this model, we simulated 10 samples with an average of 2 million paired end reads per sample, and used the isoform counts that RSEM reported to have simulated (RSEM’s ‘*sim.isoform.results*’ file) as ground truth. Isoform counts were summed to gene counts to obtain ground truth gene counts.

The Zika-infected human neuroprogenitor cell (hNPC) dataset is available at GEO database (GEO Series GSE78711). For summary statistics, we performed the analyses on all four paired end samples in the dataset and reported the mean and standard deviations across all four samples. For figures showcasing one sample, we used SRR3191542, although we performed the analysis on all four samples and found similar results across them.

### Genome and Transcriptome

We performed quantification and analysis using Ensembl *Homo sapiens* genome GRCh38 release 92 (ftp://ftp.ensembl.org/pub/release-92/fasta/homo_sapiens/dna/Homo_sapiens.GRCh38.dna_sm.toplevel.fa) and its corresponding annotation (GRCh38 release 92, ftp://ftp.ensembl.org/pub/release-92/gtf/homo_sapiens). The transcriptome was extracted from the annotation using *tophat-G*. This generation of the transcriptome file puts the genomic and pseudoaligners on an equal footing, as transcripts originating from alternate loci are not included in the transcriptome FASTA file.

### Generating TCCs

We used Salmon v0.11.2 (labeled “Salmon” or “Salmon_0.11.2” in figures) and kallisto v0.44.0 (labeled ‘kallisto’ in figures). We also included Salmon v0.8.2 in several benchmarks, which would be labeled explicitly as “Salmon_0.8.2.” Salmon and kallisto indices were built using k-mer length equal to 31. kallisto TCCs were obtained by running *kallisto pseudo*. Salmon TCCs were obtained with Salmon’s quasimapping mode by running *Salmon--dumpEQ* and reformatting Salmon’s output to match the format of TCCs in kallisto.

We used HISAT2 v2.1.0 and STAR version 2.4.2a to perform genome alignment. We used samtools v.1.2 (Li *et al.*, 2009) to sort the alignments by genomic coordinates. We then ran bam2tcc on the STAR and HISAT2 alignments to generate TCCs.

### Quantification

The TCCs generated by all four methods were quantified using kallisto’s EM algorithm, which is built on a uniform sequencing model. kallisto’s EM algorithm was run by using a branch of kallisto written specifically for this analysis (https://github.com/pachterlab/kallisto/tree/pseudoquant) and invoking *kallisto pseudoquant −l 187 −s 70* on the TCCs generated from all four methods. The *-l* and *-s* parameters correspond to the fragment size distribution (mean length and standard deviation), which are required for quantification with the EM algorithm.

### Benchmarking

In comparing the quantifications across the methods, we use the mean absolute relative distance (MARD) and the mean absolute distance. We defined mean absolute relative distance (MARD) as:

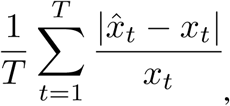

where *T* is the number of transcripts/genes considered, *x̂_t_* is the estimated quantification for transcript/gene *t*, and *xt* is the ground truth quantification for transcript/gene *t*.

We define mean absolute distance as:

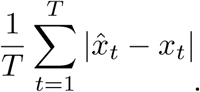

Because we use mean absolute distance on only the set of unexpressed transcripts/genes, the mean absolute distance simplifies to

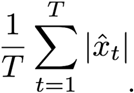

We use transcript and gene counts to calculate MARDs and mean absolute differences, obtaining gene counts from summing counts of the corresponding transcripts. We perform the Pearson and Spearman correlations on the log-transformed counts.

## Availability and Implementation

bam2tcc is available at https://github.com/pachterlab/bam2tcc. kallisto v0.44.0 containing the novel genomebam feature is available at https://pachterlab.github.io/kallisto/. The scripts and code used to regenerate our analysis are available at https://github.com/pachterlab/YLMP_2018.

## Acknowledgement

We thank Valentine Svensson for helpful feedback during the implementation of bam2tcc.

